# NF-κB/NLRP3 Translational Inhibition by Nanoligomer Therapy Mitigates Ethanol and Advanced Age-Related Neuroinflammation

**DOI:** 10.1101/2024.02.26.582114

**Authors:** Paige E. Anton, Prashant Nagpal, Julie Moreno, Matthew A. Burchill, Anushree Chatterjee, Nicolas Busquet, Michael Mesches, Elizabeth J. Kovacs, Rebecca L. McCullough

## Abstract

Binge alcohol use is increasing among aged adults (>65 years). Alcohol-related toxicity in aged adults is associated with neurodegeneration, yet the molecular underpinnings of age-related sensitivity to alcohol are not well described. Studies utilizing rodent models of neurodegenerative disease reveal heightened activation of Nuclear factor kappa-light-chain-enhancer of activated B cells (NF-κB) and Nod like receptor 3 (NLRP3) mediate microglia activation and associated neuronal injury. Our group, and others, have implicated hippocampal-resident microglia as key producers of inflammatory mediators, yet the link between inflammation and neurodegeneration has not been established in models of binge ethanol exposure and advanced age. Here, we report binge ethanol increased the proportion of NLRP3^+^ microglia in the hippocampus of aged (18-20 months) female C57BL/6N mice compared to young (3-4 months). In primary microglia, ethanol-induced expression of reactivity markers and NLRP3 inflammasome activation were more pronounced in microglia from aged mice compared to young. Making use of an NLRP3-specific inhibitor (OLT1177) and a novel brain-penetrant Nanoligomer that inhibits NF-κB and NLRP3 translation (SB_NI_112), we find ethanol-induced microglial reactivity can be attenuated by OLT1177 and SB_NI_112 in microglia from aged mice. In a model of intermittent binge ethanol exposure, SB_NI_112 prevented ethanol-mediated microglia reactivity, IL-1β production, and tau hyperphosphorylation in the hippocampus of aged mice. These data suggest early indicators of neurodegeneration occurring with advanced age and binge ethanol exposure are NF-κB- and NLRP3-dependent. Further investigation is warranted to explore the use of targeted immunosuppression via Nanoligomers to attenuate neuroinflammation after alcohol consumption in the aged.

## Introduction

The global burden of neurodegenerative diseases, including Alzheimer’s disease (AD) and AD-related dementias (ADRD), is increasing^1^. In the United States the number of people living with AD/ADRD is estimated to reach 14 million adults by 2060^1^. Advanced age (>65 years) is the leading risk factor for developing AD/ADRD^2^, and the projected increase in disease burden is thought to be attributed to the growing aging population^3^. Independent of advanced age, the specific influences that determine the onset of disease are not well understood. For example, about 90% of AD/ADRD cases are sporadic, suggesting that specific lifestyle or environmental factors may promote disease onset. Alcohol misuse has recently been identified as a potential risk factor for the development of neurodegenerative disease among the current aged population^4^. Recent surveys suggest that up to 10% of adults aged 65 years and older binge drink (having 4-5 drinks within 2 hours)^5^, and that this behavior is specifically rising in aged adult women^6^. Although alcohol misuse can cause brain damage independent of advanced age^7^, the synergistic effects of binge drinking and advanced age on the development of neurodegenerative disease are not well understood. We previously reported that intermittent binge ethanol exposure elevates neuroinflammation, early markers of neurodegeneration, and cognitive impairment in aged female mice compared to young^8^. The specific cellular and molecular mechanisms that predispose the brain of aged subjects to alcohol-related damage, however, have yet to be defined.

Exaggerated inflammatory responses of microglia (i.e. “priming”) in the brain of aged subjects is hypothesized to play a critical role in the onset of AD/ADRD^9^. Microglia, resident macrophages of the central nervous system, possess dynamic inflammatory responses that are influenced by the microenvironment. When stimulated, microglia quickly produce pro-inflammatory cytokines and enhance phagocytic processes (termed “activation”). Once the insult is removed, microglia revert from their inflammatory phenotype to a pro-resolution state and produce neurotrophic factors that support neuronal repair, including brain-derived neurotrophic factor (BDNF)^10^. Importantly, microglia lose their dynamic responses to their environment with advanced age^11^. Evidence from studies using aged rodents suggest that at baseline, microglia produce pro-inflammatory cytokines (i.e. Tumor necrosis factor α [TNF-α], interleukin 1β [IL-1β], and interleukin 6 [IL-6]) and upregulate surface activation markers like Iba-1 and CD68 compared to microglia from young counterparts ^12, 13^.

Microglia from the brains of aged mice are further characterized by a dystrophic morphology including an enlarged cell body size and shorter process length^14^. This basal inflammatory state of primed microglia can be heightened by stimuli that are common in the brains of aged subjects, including aggregated proteins and danger associated molecular patterns (DAMPs). Exposure to these factors further propagates inflammatory mediator production but ineffective phagocytosis in microglia^15–17^.

The hyperactive state of microglia in advanced aging is a contributing phenotype to CNS inflamm- aging, described by increased neuroinflammation of the brains of aged subjects compared to young at baseline^18^. The role of microglia in the onset of disease is highlighted by several studies using mouse models of neurodegeneration, in which immunosuppression of microglia with minocycline reduces neuropathologic accumulation of hyperphosphorylated tau and amyloid-beta, and neuronal apoptosis^19–21^. Clinical data also support an association between microglia priming and neurodegenerative disease development. For example, morphological characteristics of primed microglia are observed in the brains of aged humans, and single cell sequencing of microglia from aged people reveal transcriptional profiles indicative of hyperactivation^22^. Genome wide association studies have also identified several risk factors for sporadic AD development, several of which are associated with microglia pro-inflammatory signaling and impaired phagocytosis^23^. Together, these data link dysregulated immune responses of microglia to age-related neurodegenerative disease.

Although the specific factors that mediate microglia priming in advanced age are not completely elucidated, there is an emerging role of hyperactivated NLR family pyrin domain containing 3 (NLRP3) inflammasome^24, 25^. The NLRP3 inflammasome is a multimeric protein complex that requires two signals for activation. The initiating signal occurs through binding of Pattern Recognition Receptors (PRRs), leading to translocation of the nuclear factor kappa-light-chain-enhancer of activated B cells (NF-κB) to the nucleus, inducing the transcription of NLRP3-related genes (*nlrp3, pro-il1β,* and *pro- il18*). The second signal arises from intracellular stress indicators, including reactive oxygen species and/or ATP. NLRP3 then complexes with its subunit apoptosis-associated speck-like protein containing a CARD (ASC). NLRP3/ASC complexes cleave caspase-1, leading to generation of pro-inflammatory cytokines IL-1β and IL-18^25^. Rodent models of advanced age suggest an important role of NLRP3 in microglia priming. In aged mice, peripheral lipopolysaccharide (LPS) exposure induces markers of NLRP3 inflammasome activation, like IL-1β, and expression of microglia surface activation markers compared to similarly treated adult mice^26^. These data suggest hyperactivation of NLRP3 is associated with heightened microglia activation in the aged. Furthermore, aged NLRP3^-/-^ mice exhibit reduced expression of microglia activation marker Iba-1, reduced IL-1β production in the hippocampus, and improved memory behaviors compared to aged wildtype mice^13^. In rodent models of AD or tauopathy, systemic treatment with NLRP3 inhibitors reduce microglia reactivity and neuronal death in the hippocampus^27, 28^, suggesting an important role of NLRP3 in advanced age and neurogenerative diseases related neuroinflammation.

The use of systemic NLRP3 inhibitors and global NLRP3^-/-^ limit our interpretation of the role of microglia-specific NLRP3 activation in aging and neurodegeneration models. Recent evidence demonstrate genetic knockout of *nlrp3* or *asc* in microglia prevents neuronal loss and cognitive impairment in mouse models of sporadic AD^29^ and tauopathy^30^. *Ex vivo* NLRP3 inhibition prevents LPS-induced phagocytic dysfunction and pro-inflammatory cytokine production in primary microglia from young mice ^30^. Although these data suggest NLRP3 suppression is a suitable target to mitigate microglia activation and consequential neuronal damage, clinical trials utilizing NLRP3 inhibitors have variable results in alleviating inflammation in other diseases like systemic inflammatory response syndrome^31^. Thus, additional investigation into the utility of suppressing upstream activators of NLRP3, like NF-kB, is necessary. NF-κB inhibition is also shown to reduce microglia activation in mouse models of AD and in immortalized mouse microglia (BV-2s) challenged with LPS^32, 33^. Despite the relationship between NF-κB and NLRP3 activation, the effects of simultaneous inhibition of these pathways in mouse models of aging or neurodegeneration is not well defined. Taken together, these data suggest microglia-specific NF-κB signaling and downstream NLRP3 inflammasome activation may be a viable target to reduce age-related microglia dysfunction and associated neuronal damage. There remain critical gaps in knowledge that address how binge ethanol exposure may contribute to microglia inflammasome activation and associated neuronal damage in advanced age.

We, and others, have demonstrated microglia from aged mice are more reactive to ethanol compared to young^8, 34^. Yet, the mechanism leading to advance age and ethanol-induced microglial reactivity is not completely elucidated. Data from Alfonso-Loeches *et al*. suggest ethanol-related microglia reactivity in young mice is TLR4-dependent^35, 36^. Chronic ethanol feeding in young wildtype mice leads to enhanced caspase-1 expression in microglia, which is normalized in TLR4-deficient mice^36^. Because TLR4 is associated with the expression and activation of NLRP3 components, it is logical to hypothesize that ethanol contributes to TLR4-dependent activation of the NLRP3 inflammasome in microglia. Primary microglia isolated from neonatal mice respond to *ex vivo* ethanol challenge with indicators of NF-κB activation and IL-1β production, which is attenuated in microglia from TLR4^-/-^ mice^37^. Conversely, other studies fail to observe increased pro-inflammatory cytokine production in primary microglia from binge ethanol-exposed young adult male and female rats after ethanol withdrawal^38^. Together this literature highlights the necessity to characterize and compare inflammatory responses of microglia throughout the lifespan, between species, and between different ethanol exposure techniques. There is also a paucity of research exploring NF-κB and/or NLRP3- dependent responses of microglia from the brain of young or aged mice to ethanol. Despite this, the inflammasome is implicated as a key mechanistic factor leading to ethanol-related pro-inflammatory cytokine production *in vivo*. For example, young adult female mice that are chronically fed ethanol express heightened mature IL-1β in the cerebellum, which is normalized in similarly treated NLRP3^-/-^ mice. In this model, antagonism of IL-1R, the receptor for IL-1β, by anakinra treatment mitigated ethanol-induced production of pro-inflammatory and cytotoxic factors like CCL-2 and TNF-α^39^. These data, and the aging literature described above, lead us to hypothesize that NF-κB and NLPR3 inflammasome activation leads to heightened microglia reactivity and early signs of neurodegeneration in the brains of aged mice from binge ethanol exposure.

Here, we define exacerbated inflammatory responses of primary microglia isolated from the brains of aged mice to ethanol exposure *ex vivo* compared to microglia from young mice, which were mitigated by specific inhibition of the NLRP3 inflammasome. Making use of a non-toxic nanoligomer, SB_NI_112, that prevents NF-κB and NLRP3 mRNA translation^40–42^, we find that SB_NI_112 also reduced the expression of microglia activation markers in microglia from the aged mice challenged with ethanol *ex vivo*. To explore the therapeutic efficacy of SB_NI_112 *in vivo,* and because it is brain-penetrant, we employed SB_NI_112 treatment superimposed with binge ethanol exposure in aged mice. We found that SB_NI_112 treatment attenuated markers of IL-1β production, microglia reactivity, and tau hyperphosphorylation in the hippocampus of aged mice after ethanol exposure. These findings indicate NF-κB and NLRP3 may be viable therapeutic targets to normalize microglia function and prevent neurodegeneration in the current aged population that binge drinks.

## Materials & Methods

### Nanoligomer Design and Synthesis

SB_NI_112 was designed, synthesized, and provided by Sachi Bio^41–43^. SB_NI_112 is composed of an antisense peptide nucleic acid (PNA) conjugated to a gold nanoparticle. The PNA sequence is 17 base pairs long and was optimized for solubility and specificity against mRNA regions of mouse *nfkb1* (sequence: AGTGGTACCGTCTGCTA) and mouse *nlrp3* (Sequence: CTTCTACTGCTCACAGG). The PNA portion of the Nanoligomer was synthesized on a Vantage peptide synthesizer (AAPPTec, LLC) employing solid-phase Fmoc chemistry. Fmoc-PNA monomers were obtained from PolyOrg Inc. Post-synthesis, the peptides were attached to gold nanoparticles and subsequently purified using size-exclusion filtration. Measuring absorbance at 260 nm (for PNA detection) and 400 nm (for nanoparticle quantification) was performed to verify the conjugation and purity of the nanoparticle solution.

### Animals and Intermittent Binge Ethanol Protocol

All animal procedures were approved by the Institutional Animal Care and Use Committee at the University of Colorado mice were obtained from the National Institutes of Aging (NIA) aged rodent colony and maintained for at least two weeks in an in-house animal facility prior to use. To assess microglia NLRP3 expression *in vivo,* young (3-4- month-old) and aged (18–20-month-old) female C57BL/6N mice were randomized (2-3 mice/cage) into vehicle and ethanol treatment groups. Experimental cohorts were weight-matched within age groups. Mice received intragastric gavages of vehicle (water) or ethanol (3 g/kg) at 2 p.m. every-other-day over the course of 18 days, totaling 10 exposures. Eighteen hours after the final gavage, mice were anesthetized and sacrificed. Brain tissue was extracted, meninges were removed, and the left hemispheres were fixed for multi-spectral imaging. To evaluate the effect of SB_NI_112 treatment on age- and ethanol-related neurotoxicity, aged (18-20 month-old) mice were randomized (2-3 mice/cage) into vehicle and ethanol treatment groups, with or without SB_NI_112 treatment (Sachi Bio). Experimental cohorts were weight matched. Mice received intragastric gavages of vehicle (sterile water) or ethanol (3 g/kg) as described above. At the time of gavage, mice received intraperitoneal injections of SB_NI_112 (150mg/kg) or vehicle (sterile 1x PBS). Eighteen hours after the final gavage, brain tissue was harvested as aforementioned. The left hemispheres were fixed in formalin, paraffin-embedded, and sectioned for histology. Aged mice for SB_NI_112 studies were a kind gift from Drs. Elizabeth Kovacs and Travis Walrath, University of Colorado, Anschutz Medical Campus.

### Multi-Spectral Imaging

Left hemispheres from each mouse were explanted, fixed in 10% formalin and embedded with paraffin. Serial sections were cut in 10-μm-thick serial sections between bregma -2.0 to -2.2 to capture the dorsal hippocampus and adhered to a glass slide. The tissue was dewaxed with xylene, heat-treated in either pH 6 or pH 9 antigen retrieval buffer for 15 min in a pressure cooker, and blocked in antibody diluent (PerkinElmer, Waltham, MA). Sections were then sequentially stained for Iba-1 (1:500, 19741 Wako) and NLRP3 (1:50, D4D8T Cell Signaling) primary antibodies followed by HRP-conjugated secondary polymer (anti-rabbit, anti-mouse [PerkinElmer]) and HRP-reactive OPAL fluorescent reagents (PerkinElmer). To image nuclei, slides were stained with spectral DAPI and coverslips were applied with Prolong Diamond mounting media (Thermo Fisher Scientific). Whole slides were scanned using a Vectra Polaris System (Akoya Bio). After spectral unmixing, the CA3 region from each scan was manually annotated on QuPath (version 4.0) according to the Allen Mouse Brain Atlas. Qupath was used to measure the percent of Iba-1^+^ cells that also expressed NLRP3. Two CA3 sections per mouse brain was utilized for each multi-spectral imaging panel.

### Immunohistochemistry

Paraffin-embedded brains were sectioned from bregma -2.0 and -2.2 at 10- μm thickness. Sections were deparaffinized and stained with an antibody against Iba-1 (1:500, 19741 Wako), cleaved IL-1β (1:100, PA5-105048, Invitrogen), or phospho-tau-T231 (1:100, AP0035 ABclonal); slides were then counterstained with hematoxylin. Images were acquired using an upright Olympus BX43 Microscope (Waltham, Massachusetts). No specific immunostaining was seen in sections incubated with PBS/blocking buffer rather than the primary antibody (data not shown).

Staining intensities for Iba-1 and phospho-tau-T231 were measured per 10x field. Three serial sections were evaluated per mouse, and staining intensity values were averaged. All images were coded at the time of collection for a blinded analysis, and positive staining was quantified using Image J software.

### Behavioral Assays

We used then open field test to assess locomotion and anxiety-like behavior. Forty-two hours after the 7^th^ exposure, mice were placed in an open field (44cm x 44cm x 24cm) while being tracked from an overhead camera. The center and perimeter of the arena were delineated using the animal tracking software Ethovision XT (Noldus, Leesburg, VA). Time spent in the perimeter and center was determined. Anxiety-like behavior is indicated by reduced time spent in the center of the apparatus. Animals were then returned to their home cages and subjected to their 8^th^ ethanol/vehicle gavage with or without SB_NI_112 treatment (Supplemental Figure 3). To assess initial changes in spatial cognition, we then utilized a novel place preference task. Twenty-four hours after the open field task, and 18 hours after their 8^th^ exposure, animals were placed back in their open field chambers (i.e., “familiar” chamber). A door between the open field chamber and an identical “novel” chamber was opened. Animals were allowed to freely explore the familiar and novel chambers for 5 minutes. Time spent in the familiar and novel chambers was calculated using Ethovision XT (Noldus). More time spent in the novel chamber is indicative of improved spatial memory in mice.

After completion of the novel place preference task, the mice were returned to their home cages until the completion of the study.

### Primary Microglia Isolation and Culture

Primary microglia were isolated from the brains of young (3-4 months old) and aged (18-20 months old) female C57BL/6N mice as previously described^16^.

Briefly, we transcardially perfused on mice with sterile 1x PBS. Whole brain tissue was explanted and meninges were removed. Brain tissue was mechanically and enzymatically homogenized into a single cell suspension using the Miltenyi Adult Brain Isolation Kit (130-107-677) per the manufacturer’s instructions. Cell suspensions were strained with a 70 mm cell strainer and then resuspended in 6 mL of 75% stock isotonic percoll (SIP). A density gradient was then made by layering 5 mL of 35% SIP on top of the cell solution followed by 3 mL of 1x PBS. Density gradients were placed on ice for 15 minutes and then spun at *800 x g*, 4°C, for 45 minutes with slow acceleration and no break. The top 2 layers were aspirated, and microglia were collected at the 75:35 SIP interface. Cells were then washed in 45 mL of sterile 1x PBS and resuspended in Dulbecco’s modified Eagle’s medium (DMEM) (11965-092 Sigma Aldrich) supplemented with 10% fetal bovine serum and 1% penicillin-streptomycin. Immediately following isolation, primary microglia were seeded in cell culture-treated chamber slides (154941, Thermo Fisher) for immunocytochemistry or 48 well cell culture plates (130187, Thermo Fisher) for flow cytometry at a density of 3x10^5^ cells/cm^2^.

### Immunocytochemistry of Primary Microglia

Twenty-four hours after plating, cell media was replaced with serum-free DMEM overnight and then cells were challenged with 50 mM ethanol or water as vehicle. Plates were covered with Parafilm and stored at 37C, 5% CO_2_ for 24 hours.

Following ethanol or vehicle challenge, cells were fixed with 4% paraformaldehyde and quenched with 25 mM glycine in PBS. Slides were blocked for 1 hour in a solution containing 2% bovine serum albumin, 5% fish gelatin, and 0.02% saponin. Slides were stained with primary antibodies against TLR4 (1:400,1203B, RnD Systems), Iba-1 (1:200, GT10312, Invitrogen), CD68 (1:400, FA-11, BioRad), and NLRP3 (1:500, Cyro-2, Adipogen) overnight and then incubated with Alexa-fluor 488 or 568 secondary antibodies (1:500, Invitrogen) for 1 hour. Slides were washed in 1x PBS and mounted with ProLong Diamond with DAPI (P36971, Invitrogen). To evaluate NLRP3-inflammasome-dependent changes in these markers, cells were plated onto chamber slides and allowed to rest for 24 hours, and then cell media was replaced with serum free DMEM overnight. Cells were then pre-treated with 10 μM of OLT1177, 10 μM of SB_NI_112, or 1x sterile PBS as vehicle 1 hour prior to challenge with 50 mM ethanol or vehicle. 24 hours after ethanol exposure, cells were fixed and immunostained for Iba-1 and cleaved IL- 1β (1:100, PA5-105048, Invitrogen) or CD68 and cleaved caspase 1 (1:200, PA5-38099, Invitrogen) as described above. Images were acquired with a Nikon confocal microscope. Five images were acquired per chamber and the expression of each marker was determined by integrated intensity normalized to the number of DAPI-positive cells per 20x field using Image J.

### Flow Cytometry of Primary Microglia

To evaluate changes in phagocytosis caused by ethanol challenge, microglia from young and aged mice were plated and treated as described above. Twenty-four hours after ethanol challenge, ethanol-containing media was removed from all wells and cells were then treated with FluoSpheres (580/605 fluorescent, polystyrene, 1μm in diameter, F13083, Invitrogen) at a density of 200 beads/cell resuspended in serum free media for 1 hour. FluoSphere- containing media was then removed from the wells and cells were processed for flow cytometry using markers for CD11b (1:200, 101206, Biolegend) and CD45 (1:200,103132, Biolegend) to analyze the number of CD11b^+^ CD45^+^ cells that engulfed the FluoSpheres. Cytotoxicity was determined using zombie aqua following treatment with ethanol or OLT1177. Twenty-four hours after ethanol challenge, cells were harvested and stained using Zombie Aqua (1:1000, Invitrogen) for 30 minutes to assess viability. They were then blocked for Fc receptors and stained for surface markers CD11b (1:200) and CD45 (1:200) for 20 minutes. Fluorescence was determined using the CytoFlex LX Flow Cytometer (Beckman Coulter) and analysis was performed using FlowJo (version 9).

### Statistical Analysis: Values reported are means ± SEM

A total of 5 cohorts of mice were used: 3 cohorts of young and aged mice to assess microglia NLRP3 expression *in vivo*, and 2 cohorts of aged mice subjected to ethanol with and without SB_NI_112 treatment. For all *ex vivo* experiments, cells were isolated from 2-3 mice per age group, and each experiment was repeated on 2 independent occasions. The data were analyzed with GraphPad Prism (version 9.5.1) using 2-way ANOVA to compare differences between groups and Sidak’s post-hoc analysis was used to adjust for multiple comparisons. Grubb’s tests were performed on all data sets to identify statistical outliers. Statistical significance was determined by a *p-*value of 0.05 or lower. Individual *p*-values are reported in the figure legends.

## Results

### Ethanol Exposure Elevates Markers of Reactivity and NLRP3 Inflammasome Activation in Microglia of Aged Mice Compared to Young *In vivo* and *Ex vivo*

In our previously established model of intermittent binge ethanol exposure^8^, indicators of hippocampal neurodegeneration and memory impairment in aged mice are increased compared to young.

Moreover, these outcomes are associated with heightened expression of microglia reactivity markers (Iba-1, enlarged soma) and markers of NLRP3 inflammasome activation (IL-1β, NLRP3) within CA3 of the hippocampus^8^. Because microglia-specific NLRP3 activation is associated with the onset of neurodegeneration in other rodent models^29, 30^, we characterized NLRP3 expression in microglia of young and aged mice following intermittent binge ethanol exposure using multi-spectral imaging (Figure 1A). Eighteen hours after the final gavage, binge ethanol increased the proportion of microglia that expressed NLRP3 within CA3 of the hippocampus of aged mice compared to all other treatment groups (Figure 1B).

**Figure 1.**
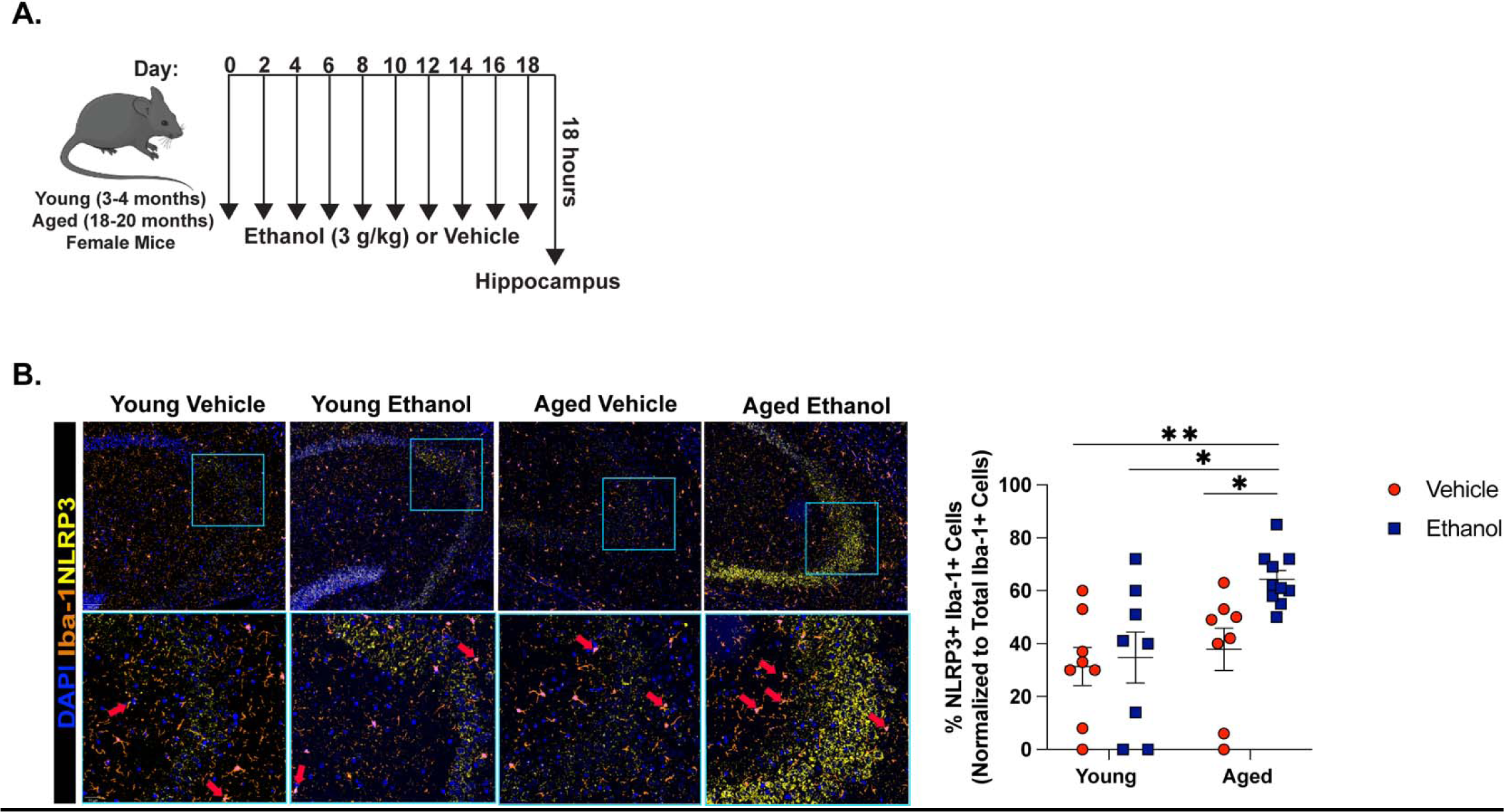
***Hippocampal Microglia from the Brains of Aged Mice Upregulate NLRP3 in Response to Binge Ethanol Compared to Young.*** (A) Young (3-4 months old) and aged (18-20 months old) female C57BL/6N mice were exposed to intragastric gavages of ethanol (3 g/kg) or vehicle every other day for 10 total exposures. Multispectral imaging was performed on left hemispheres of young and aged ethanol and vehicle treated mice. Each brain hemisphere was imaged using a whole slice scanning system and representative images at 10x and 40x magnification are shown. (B) The proportion of microglia (Iba-1^+^ cells) that were NLRP3^+^ positive in CA3 of the hippocampus was quantified using QuPath software. n = 8-10 per group, means and SEM are reported. Values were significantly different from each other determined by 2-way ANOVA with Sidak’s post hoc test, *\* p < 0.05, **p < 0.01, *** p<0.001*.

Our *in vivo* data suggest ethanol exposure elicits greater NLRP3 inflammasome activation in hippocampal microglia of aged mice compared to young. To better understand the direct effect of ethanol on NLRP3 activation in microglia, we made use of *ex vivo* primary microglia cultures. Both young and aged mice achieve blood ethanol concentrations at the level of 50 mM (230 mg/dl) or higher from our intermittent binge ethanol paradigm^8^, and blood ethanol levels are reported to match brain ethanol levels after ethanol binge in C57BL/6 mice^44^. Therefore, we treated primary microglia with or without 50 mM of ethanol. Ethanol challenge increased the expression of microglia activation marker Iba-1, but not CD68, in microglia from young mice compared to basal cells from young mice that were not ethanol treated (*p=0.05).* We did not observe any ethanol-related changes in surface markers involved in NLRP3 inflammasome activation, such as TLR4, in microglia isolated from young mice. Likewise, acute ethanol exposure did not influence expression of NLRP3 inflammasome components (NLRP3) or products (IL-1β). In microglia isolated from aged mice, ethanol caused an upregulation of Iba-1 compared to basal cells from young and aged mice (*p<0.05*), and elevated CD68 expression compared to control and ethanol-exposed microglia from young mice (*p<0.05*). In addition to elevated reactivity markers, ethanol exposure increased the expression of TLR4 in microglia from the brains of aged mice compared to basal microglia from young mice. *Ex vivo* ethanol challenge also elevated NLRP3 and IL-1β expression in these cells compared to all other treatment groups (Figure 2A-D). Given the age- and ethanol-related changes in surface markers associated with phagocytosis, like CD68, we next examined phagocytic outcomes in our *ex vivo* model. Microglia from the brains of young and aged mice were challenged with ethanol for 24 hours, and then exposed to 1μm polystyrene beads (FluoSpheres). Despite changes in CD68 expression, we did not observe any differences in the percent of cells that phagocytosed the FluoSpheres due to age or ethanol exposure (Supplemental Figure 1). These data suggest that although ethanol exposure influences expression of reactivity markers and IL-1β production in microglia from the aged mice, *ex vivo* exposure is not sufficient to impair phagocytosis of microglia from the aged brain.

**Figure 2.**
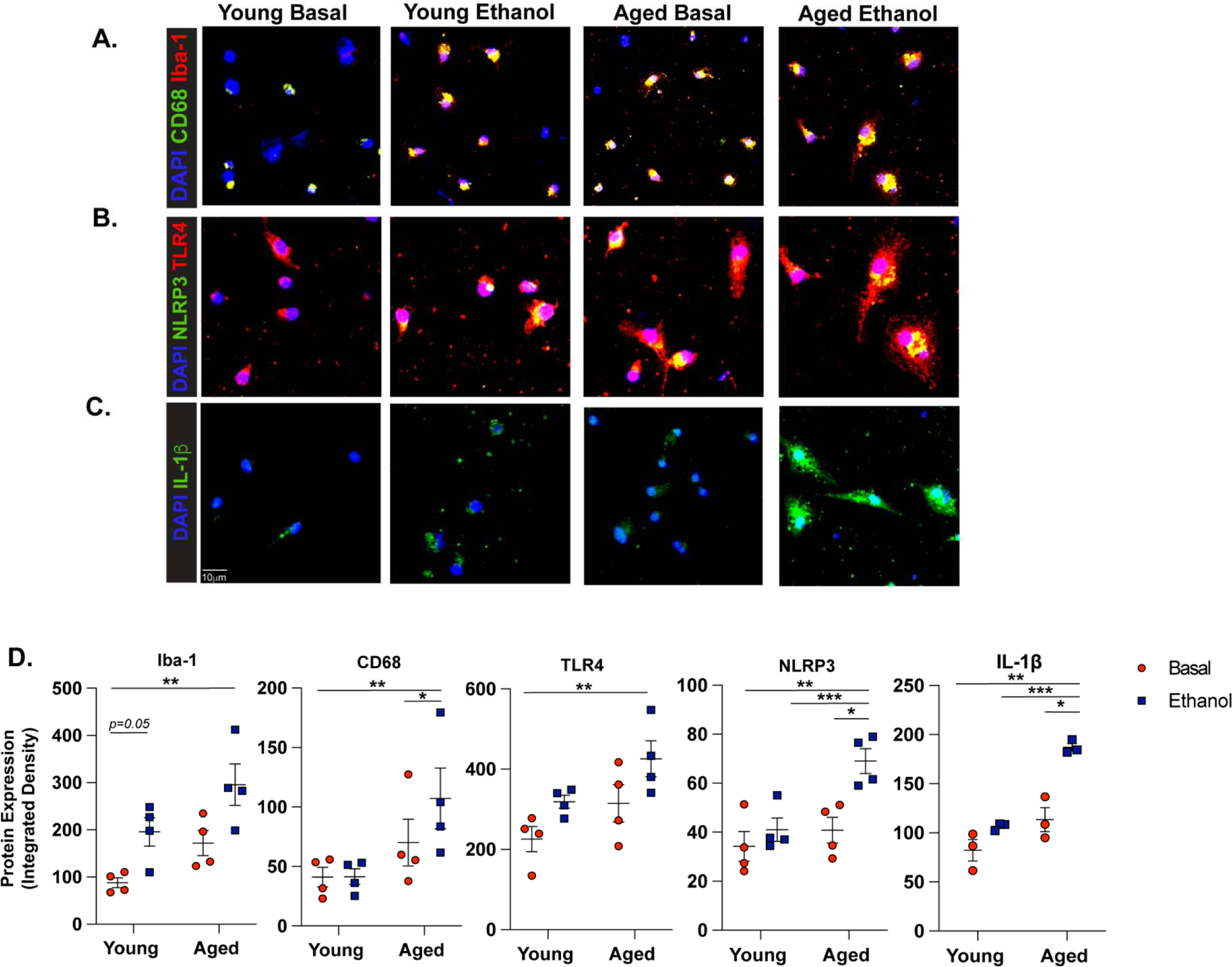
*Ex vivo Ethanol Exposure Potentiates Expression of NLRP3 and Microglia Activation Markers in Microglia from the Aged Mice Compared to Young.* Primary microglia were isolated from young and aged mice via density gradient. Cells were acclimated overnight with serum free media and challenged with or without 50 mM of ethanol for 24 hours. Cells were then fixed and processed for immunocytochemistry using antibodies against (A) CD68 and Iba1, (B) NLRP3 and TLR4, or (C) mature IL-1b. Representative images are shown at 20x objective. (D) Quantification of total staining intensity for each marker was performed using ImageJ and represented as staining intensity normalized to the number of DAPI^+^ positive cells. Individual data points are represented, n = 3-4 biological replicates per group. Means and SEM are reported, values were significantly different from each other as determined by 2-way ANOVA with Sidak’s post-hoc testing, * *p < 0.05, ** p <0.01, *** p <0.001*.

### NLRP3 Inhibition by OLT1177 and Translational Inhibition of NF-**κ**B/NLRP3 mRNA via SB_NI_112 Prevents Advanced Age- and Ethanol-Related Microglia Reactivity *Ex vivo*

Past studies show NLRP3 inhibition can attenuate microglia inflammatory responses to LPS *ex vivo,* suggesting that microglia-specific NLRP3 activation may regulate pro-inflammatory responses of microglia following ethanol exposure^27^. To investigate whether suppression of the NLRP3 inflammasome can alleviate age- and/or ethanol-related changes to microglia phenotype, we expanded our *ex vivo* ethanol exposure studies to include a small molecule inhibitor of NLRP3, OLT1177. Primary microglia were pre-treated with 10μM OLT1177 or PBS 1 hour before ethanol exposure. Immunocytochemistry revealed that OLT1177 treatment reduced ethanol-related increases in Iba-1, CD68, and IL-1β expression in primary microglia from the brains of aged mice compared to ethanol exposed, vehicle treated cells (*p<0.05*). No effect of OLT1177 on reactivity markers was observed in ethanol-exposed microglia from young mice (Figure 3A-C). Furthermore, the expression of cleaved caspase-1 in microglia from the young or aged mice was not changed following ethanol or OLT1177 treatment. Additionally, ethanol or OLT1177 treatment was not associated with cytotoxicity (Supplemental Figure 2). To explore the additional contribution of upstream activators of NLRP3, like NF-κB, to age and ethanol related phenotypes reported here, we went on to explore the effect of pre-treatment with 10 μM of SB_NI_112, a novel Nanoligomer that translationally inhibits NF-κB and NLRP3. SB_NI_112 exposure at this dose suppresses pro-inflammatory cytokine production in human derived astrocytes and human brain organoids exposed after stimulation and is not cytotoxic^41^. Like OLT177, SB_NI_112 alleviated ethanol-related increases in Iba-1, IL-1β, and CD68 (*p<0.05)* in microglia from the brains of aged mice. Interestingly, pre-treatment with SB_NI_112 elevated expression of Iba-1 in microglia from the young mice after ethanol exposure compared to vehicle treated ethanol exposed cells (*p<0.05;* Figure 4 A-C). These data suggest NF-κB signaling and NLRP3 inflammasome activation mediate enhanced IL-1β production and reactivity to acute ethanol exposure in microglia from the brains of aged mice.

**Figure 3.**
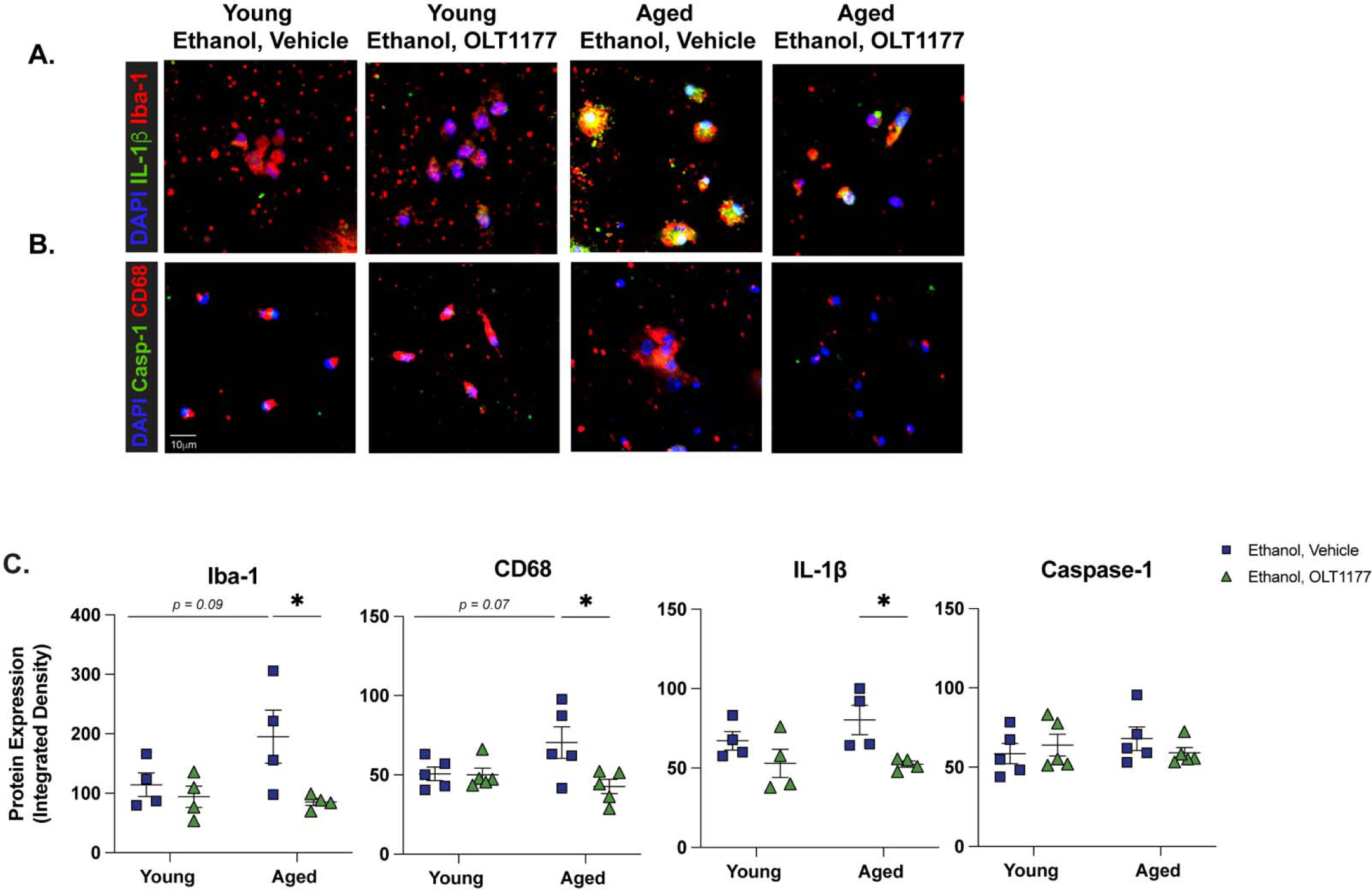
***NLRP3 Inhibition Reduces IL-1****β* ***Production and Indicators of Microglia Reactivity Due to Age and Ethanol Exposure Ex vivo.*** Primary microglia from young and aged mice were isolated via density gradient. Cells were cultured overnight with serum free media and then treated with 10μM of OLT1177 or a vehicle. One hour later, cells were challenged with 50mM ethanol. Twenty-four hours following ethanol challenge, cells were fixed and assessed for microglia reactivity markers indicators of NLRP3 inflammasome activation via immunocytochemistry. (A) Iba-1 and mature IL-1β and (B) CD68 and cleaved caspase-1. (C) Quantification of staining intensity was performed using ImageJ and represented as staining intensity normalized to the number of DAPI^+^ positive cells. Individual data points are represented, n = 4-5 biological replicates per group. Values were significantly different from each other as determined by 2-way ANOVA with Sidak’s post-hoc testing, * *p < 0.05*.

**Figure 4.**
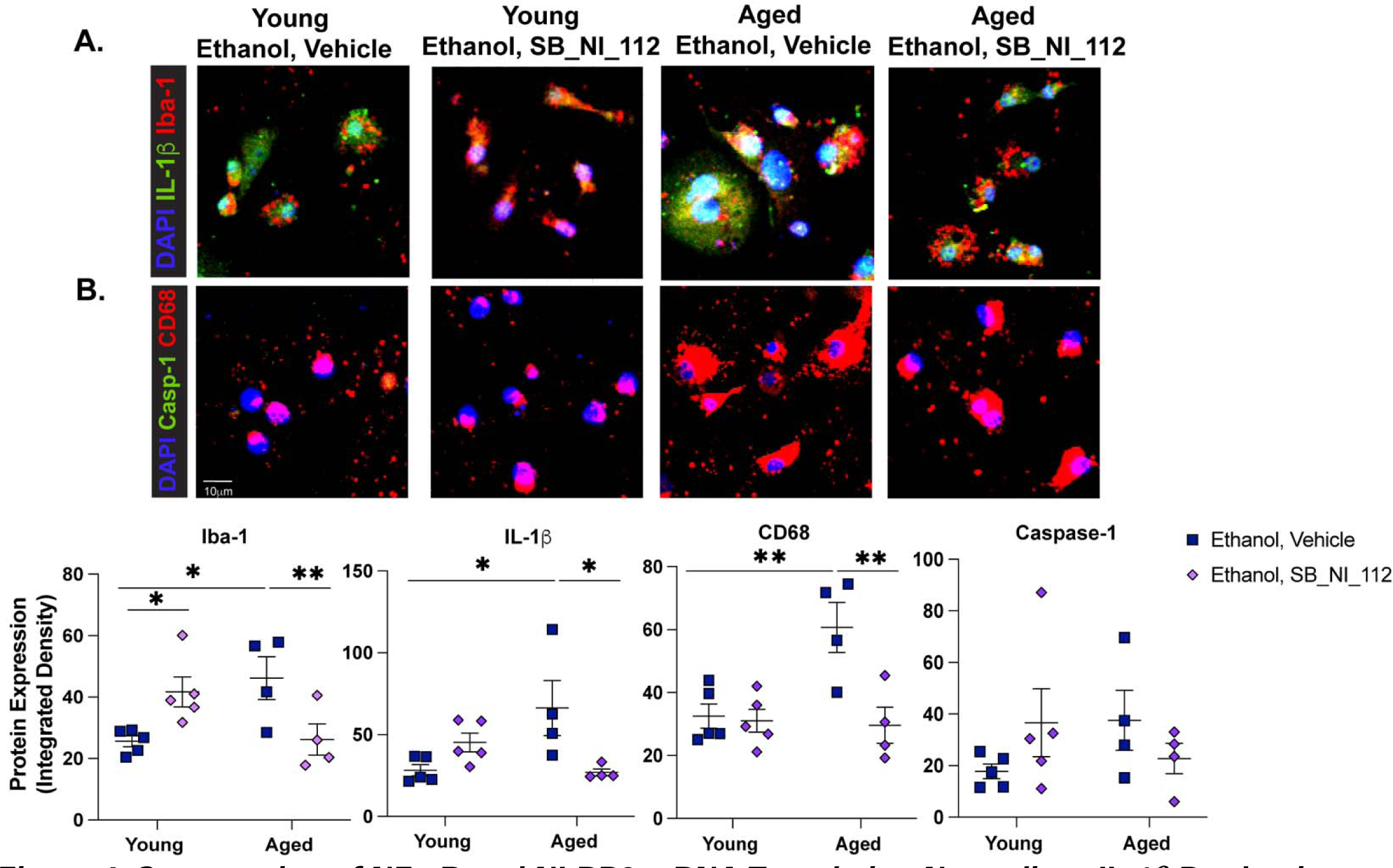
*Suppression of NF-*κ*B and NLRP3 mRNA Translation Normalizes IL-1β Production and Indicators of Microglia Reactivity Due to Age and Ethanol Exposure Ex vivo.* Primary microglia from young and aged mice were isolated via density gradient. Cells were cultured overnight with serum free media and then treated with 10μM of SB_NI_112 or a vehicle. One hour later, cells were challenged with 50mM ethanol. Twenty-four hours following ethanol challenge, cells were fixed and assessed for microglia reactivity markers indicators of NLRP3 inflammasome activation via immunocytochemistry. (A) Iba-1 and mature IL-1β and (B) CD68 and cleaved caspase-1. (C) Quantification of staining intensity was performed using ImageJ and represented as total staining intensity normalized to the number of DAPI^+^ positive cells. Individual data points are represented, n = 4-5 biological replicates per group. Values were significantly different from each other as determined by 2-way ANOVA with Sidak’s post-hoc testing, * *p < 0.05, ** p <0.01*.

Others have demonstrated that SB_NI_112 is brain-penetrant and alleviates neuroinflammation and histological features of microglia activation and tau phosphorylation in mouse models of prion disease, aging, and tauopathy^42, 45^. Importantly, SB_NI_112 reduces NF-κB and NLRP3 protein expression in the brain and is shown to be non-toxic to peripheral organs such as the liver, kidney, and colon^42, 46, 47^. To evaluate the therapeutic efficacy and potential neuroprotective role of SB_NI_112 in preventing microglia reactivity and associated neurotoxicity from ethanol in the brains of aged mice *in vivo*, we subjected aged mice to intermittent binge ethanol exposure, with or without treatment of SB_NI_112 (Figure 5A). Because NF-κB-and NLRP3 inflammasome activation is associated with IL-1β expression in the brains of mice in other models of neurodegeneration, including neurotoxin exposure ^48^ and traumatic brain injury^49^, we measured the expression of mature IL-1β in the hippocampus of aged mice after ethanol and SB_NI_112 treatment. Binge ethanol exposure increased the expression of IL-1β in aged mice compared to control (*p<0.05)*. However, SB_NI_112 treatment combined with binge ethanol exposure reduced IL-1β expression in cornu ammonis 3 (CA3) of the hippocampus of aged mice (*p<0.05;* Figure 5B-C).

**Figure 5.**
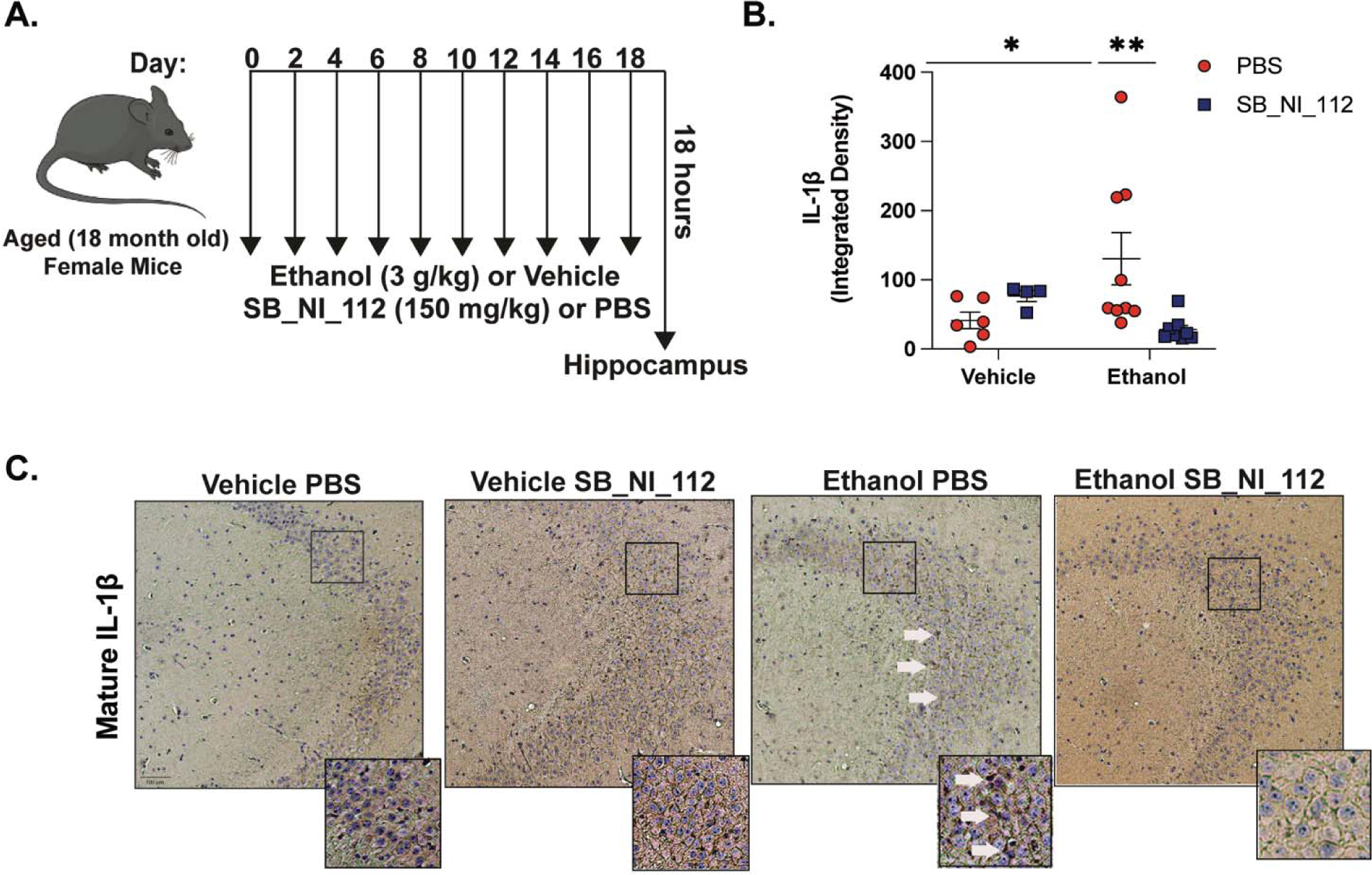
***SB_NI_112 Mitigates Advanced Age and Ethanol-Related IL-1****β* ***Production in the Hippocampus.*** (A) Aged (18-20 month-old) female mice were subjected to 10 intragastric gavages of ethanol (3g/kg) or vehicle every other day totaling 10 exposures. At the time of each gavage, animals were treated with SB_NI_112 (150mg/kg) or sterile 1x PBS via i.p. injection. Hippocampal tissue was isolated 18 hours after final exposure. (B) Paraffin-embedded mouse brain tissue was deparrafinized followed by immunohistochemistry with an antibody against mature IL-1β . (C) Expression of IL-1β staining in CA3 of the hippocampus from all treatment groups shown at 10x magnification. Insets show neuronal ribbon at 20x magnification, white arrow indicates IL-1β staining. Individual staining intensity values are represented, n = 4-9 per group. Values were significantly different from each other determined by 2-way ANOVA with Sidak’s post hoc test, * *p < 0.05, ** p <0.01*.

Microglia are an important source of pro-inflammatory cytokines in the brains of young mice after ethanol exposure, as demonstrated by microglia depletion studies^50^. Given these previous reports, we evaluated the effects of SB_NI_112 on microglia phenotype in aged mice after binge ethanol. Ethanol exposure increased the expression of cell-surface microglia reactivity marker Iba-1 in CA3 of aged mice compared to aged controls (*p<0.05)*. Microglia within CA3 also exhibited morphological features consistent with pro-inflammatory polarization due to ethanol exposure compared to control, such as an enlarged soma size (Figure 6A). SB_NI_112 attenuated ethanol-related inductions in Iba-1 and microglia soma size compared to aged animals challenged with binge ethanol alone (Figure 6 A,B).

**Figure 6.**
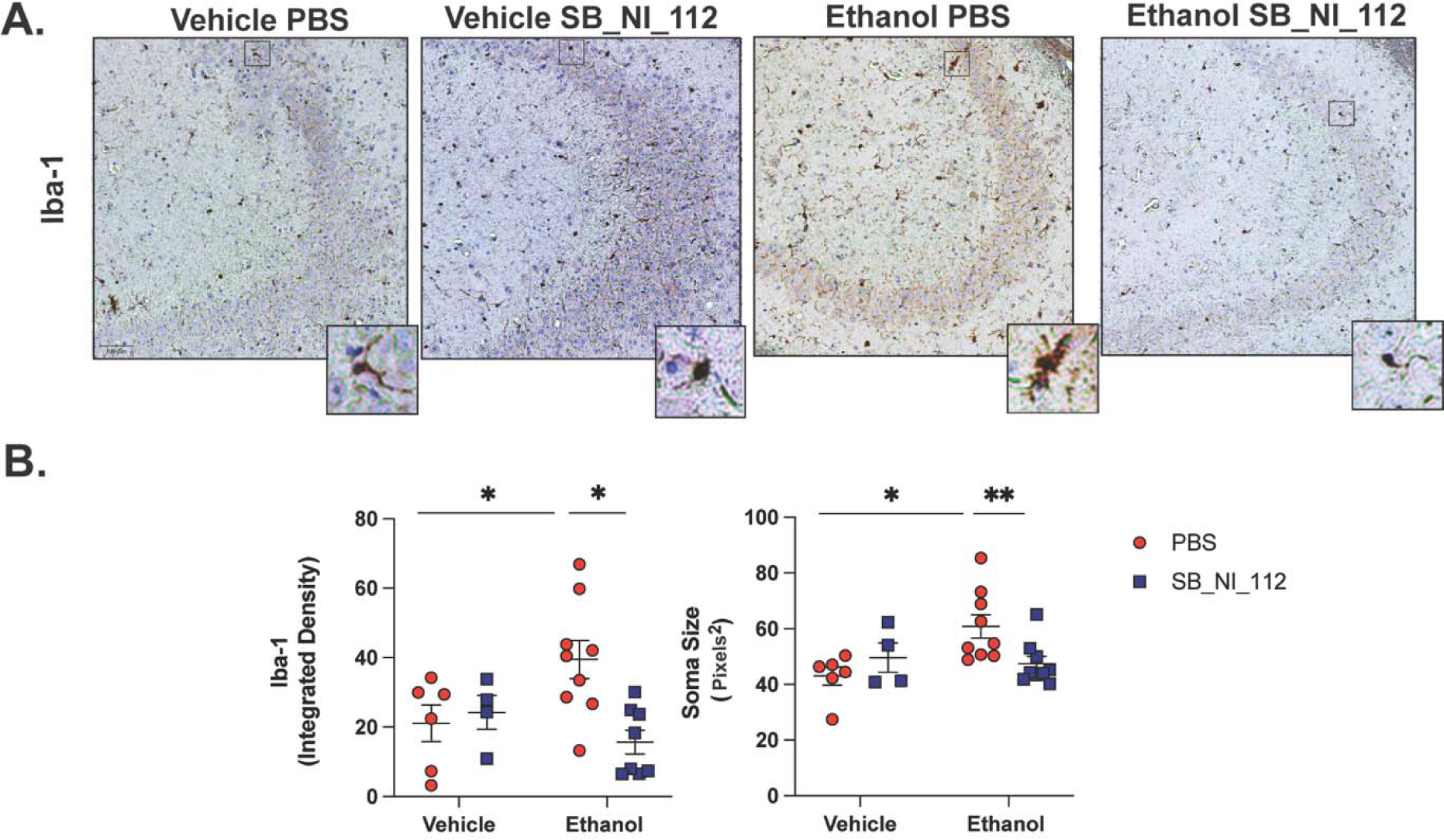
*Ethanol Related Increases in Microglia Reactivity Markers are Reduced by NF-*κ*B and NLRP3 mRNA Suppression in the Hippocampus of Aged Mice.* Aged female mice were subjected to our intermittent binge ethanol exposure paradigm, consisting of 3g/kg ethanol or vehicle gavages every-other day, totaling 10 exposures. Animals were treated with i.p. injections of SB_NI_112 (150mg/kg) or sterile 1x PBS at the time of gavage. (A) Paraffin-embedded mouse brain tissue was deparrafinized followed by immunohistochemistry with an antibody against microglia reactivity marker Iba-1. Expression of Iba-1 staining in CA3 of the hippocampus from all treatment groups with representative images of microglia morphology shown to the right of each image at 20x magnification. (B) Quantification of Iba-1 staining intensity and average microglia soma area. Individual values are represented, n = 4-9 per group. Values were significantly different from each other determined by 2- way ANOVA with Sidak’s post hoc test, * *p < 0.05 ** p < 0.01*.

These data suggest NF-κB and NLRP3 are important mediators of age and ethanol-related microglia reactivity in the hippocampus.

Microglia are implicated to play an intimate role in regulating tau hyperphosphorylation and neuronal death in neurodegenerative disease^51^. Elevated expression of IL-1β has been previously associated with neuronal death and tau hyperphosphorylation in models of ethanol expsoure^52^ as well as in transgenic mouse models of AD^53^. Because SB_NI_112 suppressed ethanol-related hippocampal IL- 1β expression and markers of microglia reactivity, we next evaluated outcomes of neurodegeneration, including the expression of tau phosphorylation at pathogenic site T231. Binge ethanol increased the expression of T231 in CA3 of aged mice compared to aged control animals (*p<0.05).* SB_NI_112 protected against ethanol-induced tau hyperphosphorylation, as ethanol and SB_NI_112 animals had reduced expression of T231 in CA3 compared to ethanol vehicle exposed animals (*p<0.05)*. There was no difference in T231 expression between control mice and mice given SB_NI_112 alone (Figure 7). Because tau hyperphosphorylation and expression of neuroinflammatory markers are associated with neuronal apoptosis, we performed TUNEL staining in the hippocampi of aged, ethanol, and SB_NI_112 exposed animals. SB_NI_112 did not impact the number of TUNEL^+^ cells in control or ethanol treated mice in CA3 of the hippocampus (data not shown). We have previously shown that intermittent binge ethanol impairs contextual memory in aged mice 3 days after final exposure^8^. To test if SB_NI_112 impacts hippocampal-related cognition of aged and ethanol exposed mice we investigated behavioral outcomes in the open field task and novel place preference task (Supplemental Figure 3). The open field task was employed after the 7^th^ gavage and i.p injection. We did not observe any ethanol- or Nanoligomer-related changes in anxiety-like behavior between treatment groups. Mice also did not exhibit differences in spatial memory via the novel place preference task 18 hours after the 8^th^ exposure (Supplemental Figure 3). Although SB_NI_112 was effective in mitigating neuroinflammatory outcomes in the brains of aged mice after 18 hours binge ethanol exposure, we were limited to observe ethanol-or Nanoligomer-related differences on behavioral assays throughout the course of our binge ethanol and SB_NI_112 exposure paradigm.

**Figure 7.**
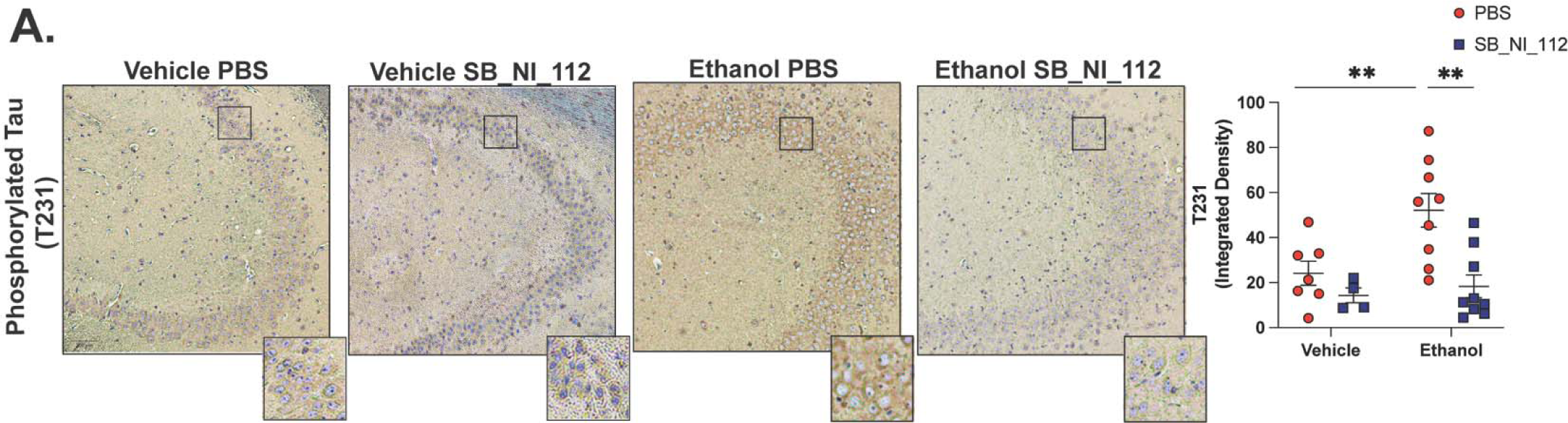
***Hippocampal Tau Hyperphosphorylation Due to Advanced Age and Ethanol Exposure are Alleviated by SB_NI_112.*** Aged female mice were exposed to 10 gavages of 3g/kg ethanol or vehicle every-other day, with or without SB_NI_112 treatment. Paraffin-embedded mouse brain tissue was deparrafinized followed by immunodetection (A) tau phosphorylation site T231. Counterstaining was performed to visualize nuclei with hematoxylin. Individual values are represented, n = 4-9 per group. Values were significantly different from each other as determined by 2-way ANOVA with Sidak’s post-hoc testing, *\*\* p <0.01*.

## Discussion

Despite the increasing incidence of binge drinking among older Americans ^54^, relatively little is known about the molecular mechanisms that predispose the brains of these aging individuals to alcohol-related neuroinflammation and damage. Here, we report that intermittent binge ethanol exposure in aged mice leads to increased NLRP3 expression in hippocampal microglia compared to young *in vivo*. Similarly, we observed that primary microglia isolated from aged mice express increased activation markers 24 hours after ethanol exposure *ex vivo* compared to primary microglia from the brains of young mice. Age- and ethanol-related increases in these outcomes were mitigated by an NLRP3- specific inhibitor and translational inhibition of NF-κB and NLRP3 via SB_NI_112. These data led us to investigate the effects of SB_NI_112 on ethanol-related neuroinflammation in our ethanol exposure and advanced age model. We found SB_NI_112 reduced microglia reactivity and IL-1β expression in the hippocampus of aged mice. Importantly, decreased neuroinflammatory markers via SB_NI_112 treatment were associated with reductions in early signs of hippocampal neurodegeneration, such as tau hyperphosphorylation.

Our *in vivo* data show that a greater proportion of microglia express NLRP3 after binge ethanol in the hippocampus of aged mice compared to young. These data build on past reports that ethanol exposure promotes NLRP3 activation in microglia of young mice^36^, and that microglia become more reactive to ethanol in the brains of aged mice^8, 34^. It is currently unclear if pro-inflammatory phenotypes of microglia in the young or aged brain are a direct consequence of ethanol exposure, a result of DAMPs generated due to ethanol neurotoxicity, or both. For example, primary microglia from the young mouse brain exposed to ethanol *ex vivo* increase the expression of TLR4 and Iba-1, NF-κB signaling, IL-1β production^37^, and reactive oxygen species generation^36^. These reports suggest ethanol is sufficient to induce microglia pro-inflammatory polarization and NLRP3 inflammasome activation. Some data support the hypothesis that ethanol can promote TLR4/TLR2 dimerization through disruptions in lipid rafts in microglia, initiating downstream signaling in the absence of a ligand^55^. Conversely, evidence from rodent brain slice cultures exposed to ethanol suggest DAMPs produced after ethanol challenge, like HMGB1, are necessary for the inflammatory effects of ethanol^56^.

To begin to understand the age dependent effects of direct ethanol exposure, we utilized primary microglia cultures from young and aged mice. For the first time, we show that primary microglia from the brains of aged mice upregulate surface activation markers (Iba-1, CD68, TLR4) and indicators of inflammasome activation (NLRP3, IL-1β) 24 hours after acute ethanol exposure *ex vivo* compared to young. These data corroborate past reports that ethanol directly induces microglia reactivity^57^.

However, microglia are found to produce TLR4-ligands, like HMGB1, in response to ethanol in rat brain slice cultures^58^. Thus, future investigation utilizing HMGB1 antagonists in young or aged mice *in vivo* and *ex vivo* may be necessary to fully differentiate the contribution of ethanol and DAMPs to neuroinflammation aged mice.

Increased expression of surface activation markers and NLRP3 inflammasome markers in microglia from the brains of aged mice in response to ethanol support other studies that show primary microglia from aged rats^59^ and mice^16^ increase surface activation markers and produce more IL-1β after LPS exposure compared to cells isolated from their younger counter parts. NF-κB signaling and NLRP3 inflammasome activation are reported to regulate pro-inflammatory responses of microglia from the young brain^29, 60^, but role of NF-κB and/or NLRP3 in regulating activation of primary microglia from the brain of aged subjects, with or without an inflammatory stimulus like ethanol, is not well described. Here, we report that specific inhibition of NLRP3 or suppression of NF-κB and NLRP3 mRNA via SB_NI_112 attenuates ethanol-related increases in Iba-1, CD68, and IL-1β in primary microglia from aged mice. Others have shown NLRP3 activation in microglia from neurodegeneration models is associated with decreased phagocytosis^61^ and acute ethanol exposure in primary rat microglia suppresses amyloid beta phagocytosis^62^. However, others studies suggest acute ethanol exposure in primary mouse microglia increases phagocytosis^37^. To explore age and ethanol-dependent effects in phagocytosis in our culture system, we exposed primary microglia to FluoSpheres after ethanol treatment. However, we did not see any changes in the phagocytic efficacy of these cells. Together, these data suggest that NLRP3 is an important mediator of initial changes in microglia reactivity to ethanol occurring with advanced age.

The molecular mechanism leading to NLRP3 inflammasome dysregulation in microglia from the brains of aged mice is not fully elucidated. Here, we show that primary microglia from aged mice respond to acute ethanol exposure with an induction in TLR4 expression over young. Given the important role of TLR4 in ethanol-related microglia reactivity in the young mice^36, 37, 63^, increased TLR4 expression and binding may be an avenue by which microglia from the aged brain are susceptible to NLRP3 inflammasome activation from ethanol. We also show microglia produce more IL-1β with advanced age and ethanol exposure. IL-1β can bind to its receptor, IL-1R, and initiate NF-κB signaling and contribute NLRP3 activation^53^. Therefore, enhanced IL-1β production may sustain NLRP3 activation in primary microglia from aged mice after ethanol exposure. Additionally, others have shown *in vitro* microglia aging systems, like high-division immortalized microglia, produce more reactive oxygen species (ROS) in response to beta-amyloid stimulation^64^. Increased microglial ROS generation from ethanol exposure may contribute to inflammasome signaling. The use of TLR4 and IL-1R antagonists or antioxidant treatments in microglia culture systems of advanced age will help to fully define NLRP3 dysregulation in microglia from the brains of aged subjects.

We did not observe a strong ethanol-related phenotype in primary microglia from young mice, unlike past reports. These discrepancies may be due to differences in ethanol dose, as these past reports utilize up to 100 mM of ethanol in an acute exposure^50, 65^. Here, microglia were isolated from mice at 3-4 months of age, while others utilize primary microglia from neonatal mice, which may be more vulnerable to ethanol-related toxicity^37^. Our work contradicts studies that suggest ethanol suppresses pro-inflammatory signaling in immortalized rat microglia after acute ethanol exposure^66^. Our data and past literature underscore the need for further investigation into the effects of ethanol on microglia function between different species and age groups. Interestingly, we observed that pre-treatment with SB_NI_112, but not OLT1177, before ethanol exposure increased Iba-1 expression in microglia from young mice *ex vivo*. Although NF-κB promotes the expression of pro-inflammatory cytokine mRNA, NF-κB can also negatively regulate pro-inflammatory responses over time^67^. Genetic deletion of NF-κB subunit, p50, promotes pro-inflammatory responses of microglia from young mice challenged with LPS^68^. These data suggest that translational inhibition NF-κB mRNA in microglia from the brains of young mice exposed to ethanol may have mitigated the immuno-regulatory role of NF-κB. To better understand the dynamic effects of NF-κB translational inhibition, future studies should employ time course studies to evaluate microglia activation markers due to ethanol and/or SB_NI_112 exposure. Further investigation into the role of NF-κB in regulating microglia activation states throughout the lifespan or in the context of ethanol exposure will lead to a better understanding on when, and if, targeting NF-κB signaling is an effective therapeutic target for neuroinflammation and resulting injury.

Given the anti-inflammatory effect of SB_NI_112 administration in primary microglia from aged mice after ethanol exposure, we investigated the efficacy of SB_NI_112 treatment in alleviating binge ethanol and advanced age neuroinflammation *in vivo*. Similar to our past findings, we show that intermittent binge ethanol exposure increases expression of mature IL-1β in CA3 of the hippocampus, as well as promotes Iba-1 expression and markers of microglia dystrophy, such as enlarged soma size^8^. Ethanol-related outcomes were attenuated by SB_NI_112 administration at the time of ethanol gavage, suggesting a role for NF-κB signaling and NLRP3 inflammasome activation in promoting neuroinflammation in this model. Normalizing microglia dysfunction and NLRP3 inflammasome activation ameliorate tau hyperphosphorylation in the hippocampus of transgenic mice^28, 42, 45^.

Therefore, we investigated the therapeutic efficacy of SB_NI_112 in preventing tau phosphorylation in the hippocampus of aged mice exposed to binge ethanol. Ethanol exposure increased expression of pathological tau phosphorylation site T231 in aged mice compared to controls, which was reduced with SB_NI_112 treatment at the time of ethanol exposure. These data corroborate evidence suggesting SB_NI_112 is effective at preventing tau phosphorylation in other models of tauopathy^45^.

Microglia are considered potent sources of IL-1β production in rodent models of advanced age or ethanol exposure. Specific mitigation of the NLRP3 inflammasome in microglia alleviates neurodegenerative disease phenotypes in rodents^29, 30^. This literature, with our current data, may suggest NLRP3 inflammasome activation in microglia promotes IL-1β generation and pathological phosphorylation of tau in the hippocampus of ethanol exposed aged mice. However, both neurons and astrocytes are also shown to upregulate NLRP3 in response to ethanol in young mice^69–71^.

Therefore, future studies utilizing specific knockout of NLRP3 in microglia are needed to determine the role of microglia NLRP3 inflammasome activation in this model.

We explored initial changes in cognition due to ethanol exposure and/or SB_NI_112 treatment in our aged mice before tissue harvest. We did not observe any differences in behavior on the open field task or any changes in place preference in the novel place preference task due to ethanol or SB_NI_112 treatment. Our previous reports show that binge ethanol exposure impairs cognition in aged mice^8^. However, those behavioral assays were done 2 and 3 days after final ethanol exposure and separate cohorts of mice were used for tissue collection and behavioral assays. Other models of ethanol-related cognitive impairment using young rodents also utilize behavioral assays several days after ethanol consumption to eliminate confounding effects of ethanol withdrawal^72^. Given the limited number of animals available to perform aging studies, the use of behavioral assays throughout the ethanol exposure paradigm is a limitation of the current study. Future studies should utilize our model of advanced age and binge ethanol with SB_NI_112 treatment and behavioral assays several days after ethanol exposure to fully determine the effects on cognition. Despite elevated tau phosphorylation in the hippocampus of ethanol-exposed mice, we failed to observe differences in neuronal apoptosis, and thus suggest the ethanol-related phenotypes reported here are indicative of early changes in neurodegenerative disease phenotypes that may precede neuronal death. Lastly, our current study did not address changes in peripheral factors that may influence neuroinflammation in this model. For example, advanced age and ethanol are independently shown to increase circulating LPS^73, 74^, which can contribute to NF-κB mediated NLRP3 inflammasome activation in the brain^75^. Because both ethanol and SB_NI_112 are administered systemically, additional investigation into the contribution of peripheral inflammation to this model of neuroinflammation is necessary.

In conclusion, our data implicate NF-κB and the NLRP3 inflammasome in ethanol-related neuroinflammation and markers of neurodegeneration occurring with advanced age. We show that inhibition of NF-κB and NLRP3 mRNA translation alleviates activation of microglia from ethanol *ex vivo* and *in vivo* and mitigates ethanol-related neuropathology in the brains of aged mice. These findings add to the growing literature that Nanoligomers may be safe and effective therapeutics to treat age-related neurodegeneration^41, 45–47^. Our studies warrant additional investigation into the use of Nanoligomers for targeted immunosuppression of NF-κB and NLRP3 to slow the onset of neurodegeneration in the current aged population that binge drinks.

## CRediT author statement

Paige Anton: Conceptualization, Validation, Formal analysis, Investigation, Writing – Original Draft, Writing – Review & Editing; Prashant Nagpal, PhD: Conceptualization, Validation, Formal analysis, Writing – Review & Editing; Julie Moreno, PhD: Conceptualization, Validation; Matthew A. Burchill, PhD: Formal analysis, Writing – Review & Editing; Anushree Chatterjee: Formal Analysis; Nicolas Busquet, PhD: Formal analysis, Writing – Review & Editing; Michael Mesches, PhD: Formal analysis; Elizabeth J. Kovacs: Writing – Review & Editing, Visualization, Supervision, Project administration, Funding acquisition; Rebecca McCullough, PhD: Conceptualization, Validation, Formal analysis, Investigation, Writing – Review & Editing, Visualization, Supervision, Project administration, Funding acquisition

