## Supplementary Figures for "NF-κB/NLRP3 Translational Inhibition by Nanoligomer Therapy Mitigates Ethanol and Advanced Age-Related Neuroinflammation"

**Supplementary Figure**

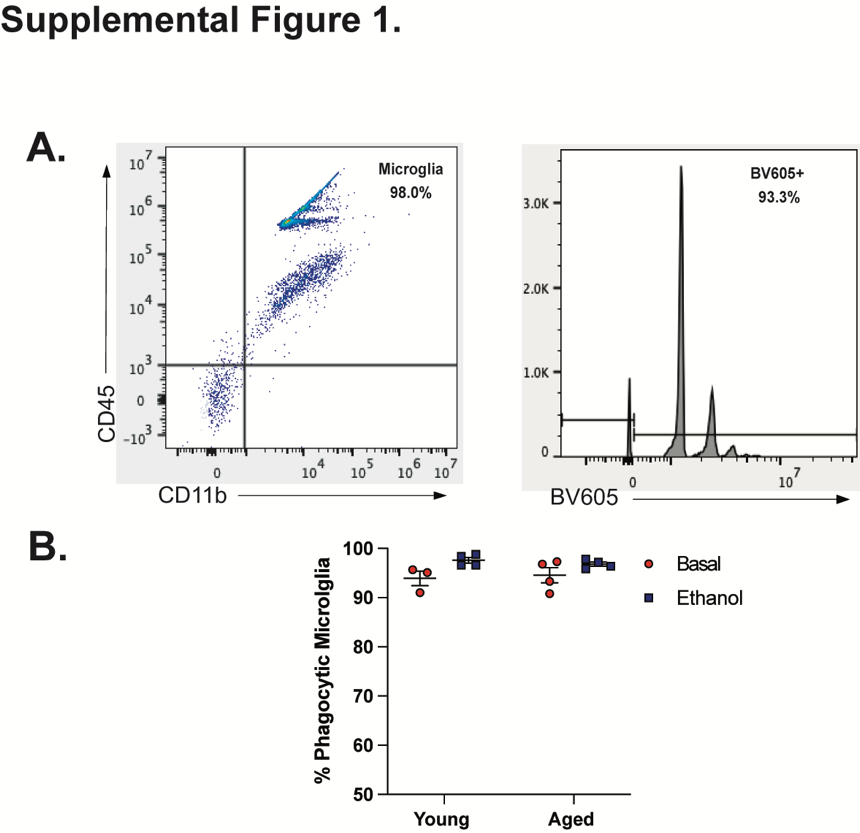

***Supplemental Figure 1. No Differences in Phagocytosis in Primary Microglia from the Young or Aged Mouse Brain After Ethanol Exposure.*** Cells were plated and challenged with 50 mM ethanol or vehicle for 24 hours. Ethanol-containing media was then removed from cells and replaced with Fluosphere (580/605 fluorescent) containing media for 1 hour. Percent phagocytic cells were then measured via flow cytometry. (A) Representative gating strategy of the percent of phagocytic microglia (CD11b^+^CD45^+^ cells that were also 605 positive). Representative graphs taken from a young, basal sample. (B) Quantification of the percent phagocytic microglia from each group, normalized to total CD11b^+^CD45^+^ cells.

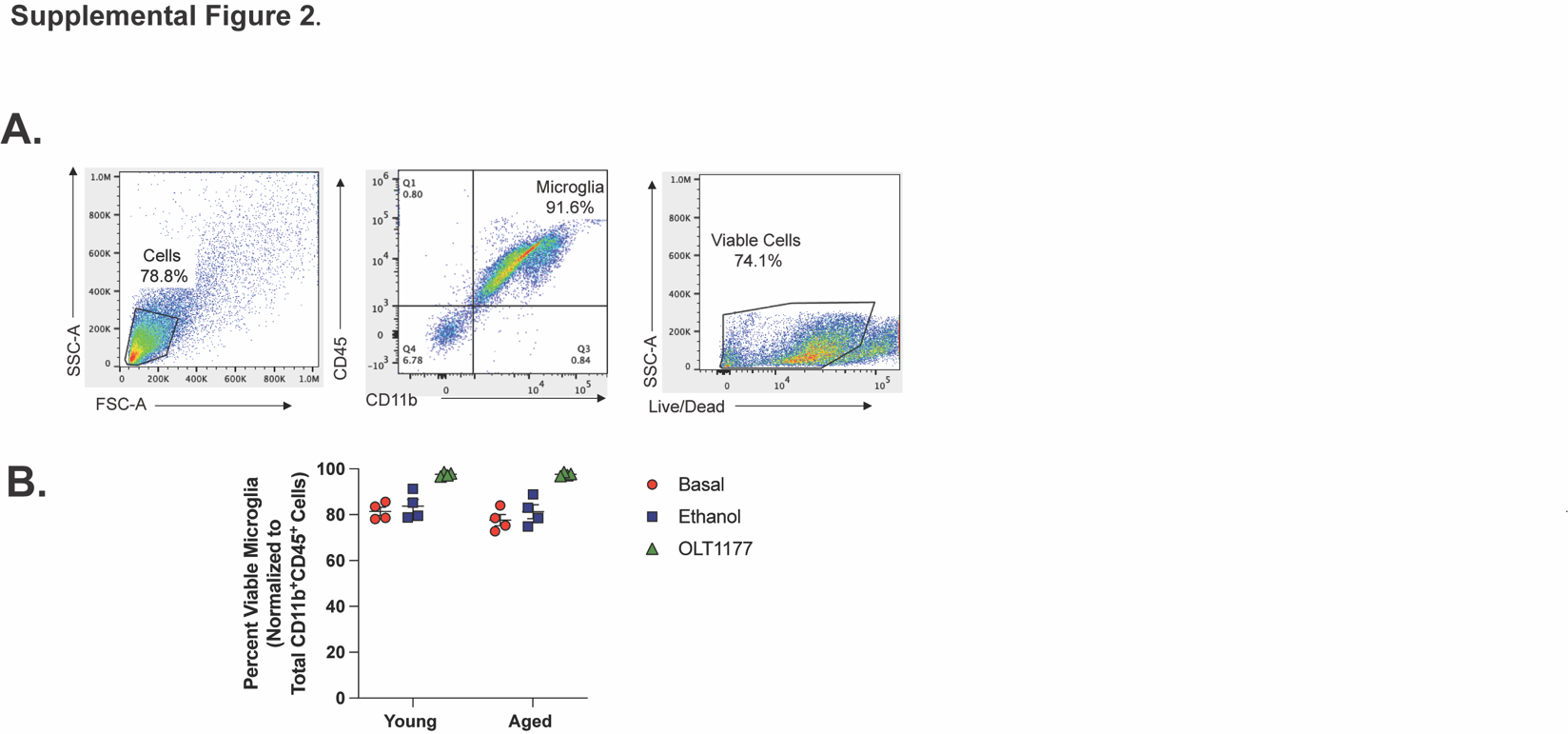

***Supplemental Figure 2.* Ex Vivo *Ethanol or OLT1177 Treatment Did Not Change Cell Viability.***

Cells were plated challenged with 50 mM ethanol or 10 μM OLT1177 for 24 hours and then processed for flow cytometry to assess microglia enrichment and cell death with specific macrophage (CD11b and CD45) and viability (Zombie Aqua) markers. (A) Gating strategy to capture percentage of viable microglia. Representative image taken from a young, basal sample. (B) Percent viable microglia, normalized to the total number of CD11b^+^CD45^+^ cells. Individual data points are represented, n = 3-4 biological replicates per group.

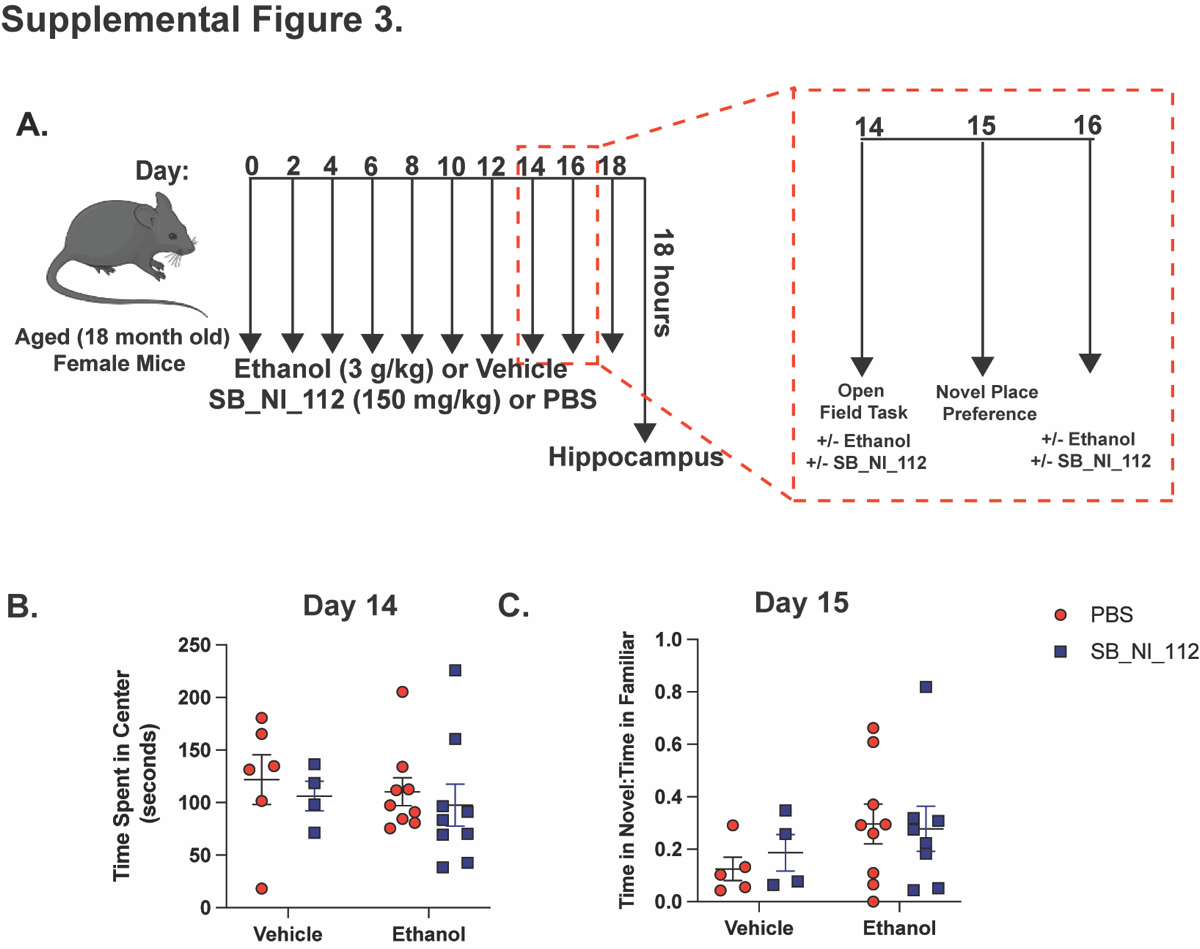

***Supplemental Figure 3. No Differences in Anxiety-Like Behavior or Place Preference Due to Binge Ethanol Exposure and/or SB_NI_112 Treatment in Aged Mice.*** Aged female mice were subjected to our intermittent binge ethanol exposure paradigm, consisting of 3g/kg ethanol or vehicle gavages every-other day, totaling 10 exposures. Animals were treated with i.p. injections of SB_NI_112 (150mg/kg) or sterile 1x PBS at the time of gavage. (A) Behavior testing timeline. All animals were subjected to the Open Field Task 42 hours after their 7^th^ exposure (Day 14). Animals participated in the Novel Place Preference Task 18 hours after their 8^th^ exposure (Day 15). They were then returned to their home cages and until the completion of the study. (B) Total time spent in the center of the open field arena in seconds. Animals were placed in the open field arena for 10 minutes. (C) Ratio of time spent in the novel chamber compared to familiar chamber on Day 15. Animals were subjected to the Novel Place Preference Task for 5 minutes.
